# Comprehensive statistical inference of the clonal structure of cancer from multiple biopsies

**DOI:** 10.1101/125138

**Authors:** Jie Liu, John T. Halloran, Jeffrey A. Bilmes, Riza M. Daza, Choli Lee, Elisabeth M. Mahen, Donna Prunkard, Chaozhong Song, Sibel Blau, Michael O. Dorschner, Vijayakrishna K. Gadi, Jay Shendure, C. Anthony Blau, William S. Noble

**Author notes:** Correspondence to Dr. C. Anthony Blau, 850 Republican St., Box 358056, University of Washington, Seattle, WA 98109, USA and Dr. William S. Noble, 3720 15th Ave NE, Box 355065, University of Washington, Seattle, WA 98195, USA.

## Abstract

A comprehensive characterization of tumor genetic heterogeneity is critical for understanding how cancers evolve and escape treatment. Although many algorithms have been developed for capturing tumor heterogeneity, they are designed for analyzing either a single type of genomic aberration or individual biopsies. Here we present THEMIS (Tumor Heterogeneity Extensible Modeling via an Integrative System), which allows for the joint analysis of different types of genomic aberrations from multiple biopsies taken from the same patient, using a dynamic graphical model. Simulation experiments demonstrate higher accuracy of THEMIS over its ancestor, TITAN. The heterogeneity analysis results from THEMIS are validated with single cell DNA sequencing from a clinical tumor biopsy. When THEMIS is used to analyze tumor heterogeneity among multiple biopsies from the same patient, it helps to reveal the mutation accumulation history, track cancer progression, and identify the mutations related to treatment resistance. We implement our model via an extensible modeling platform, which makes our approach open, reproducible, and easy for others to extend.

---

Cancer is heterogeneous in the sense that the cancer cells in a tumor are not genetically identical, but form distinct clones, defined as subpopulations of cancer cells that host the same genomic aberrations. In aggressive and metastatic cancers, these genomic aberrations quickly evolve, resulting in extreme spatial and temporal heterogeneity [1, 2]. Therefore, multiple biopsies over different locations and at different time points need to be collected and sequenced in order to capture the complexity of tumor genomic landscapes and provide insight into how tumors evolve and escape treatment [3, 4]. Accordingly, computational tools are needed to accurately characterize the clonal structure of cancer and reveal how that structure evolves over time.

In recent years, a large number of computational tools and statistical models have been developed to analyze tumor heterogeneity from DNA sequencing data (**Table 1**). However, most of these tools only model one type of genomic aberration, such as single-nucleotide variants (SNVs), copy number alterations (CNAs), or insertions and deletions (indels). Restricting the analysis to a single type of mutation not only reduces statistical power to accurately detect the clonal structure within the tumor, but also prevents us from understanding interactions among different types of mutations. Furthermore, many SNV-based methods assume that no copy number changes have occurred, which is extremely improbable. Therefore, their estimation of the prevalence of a given clone can be inaccurate, and the corresponding heterogeneity results may be misleading. Existing methods that capture SNVs and CNAs in the same model (i.e., phyloWGS [5], SPRUCE [6] and Canopy [7]) require running a CNA-calling algorithm before heterogeneity analysis, but accurate CNA characterization also depends on heterogeneity analysis.

**Table 1:** Software tools for characterizing within-patient and intra-tumor heterogeneity and their features, including whether they capture SNVs, CNAs and indels, whether they support multiple biopsy analysis, and their key models and algorithms.

| Software | Year | SNV | CNA | Indel | Multiple | Model/Algorithm |
| --- | --- | --- | --- | --- | --- | --- |
| OncoSNP [8] | 2010 |  | ✓ |  |  | Mixture model, EM, Bayesian methods |
| TuMult [9] | 2010 |  | ✓ |  | ✓ | Breakpoint distance |
| GRAFT [10] | 2012 |  | ✓ |  |  | Partial maximum likelihood |
| ABSOLUTE [11] | 2012 | ✓ | ✓ |  |  | Maximum likelihood |
| TrAp [12] | 2013 | ✓ |  |  |  | Exhaustive search under constraints |
| THetA [13, 14] | 2013 |  | ✓ |  |  | Maximum likelihood |
| CancerTiming [15] | 2013 |  | ✓ |  |  | Maximum likelihood |
| OncoSNP-seq [16] | 2013 |  | ✓ |  |  | Mixture model, EM, Bayesian methods |
| PyClone [17] | 2014 | ✓ |  |  | ✓ | Dirichlet Process, beta-binomial/MCMC |
| SciClone [18] | 2014 | ✓ |  |  | ✓ | Beta mixture model/variational Bayes |
| Clomial [19] | 2014 | ✓ |  |  | ✓ | Binomial mixture model, EM |
| CloneHD [20] | 2014 | ✓ | ✓ |  | ✓ | HMM, EM, variational Bayes |
| MEDICC [21] | 2014 |  | ✓ |  | ✓ | Finite state transducer, minimum-event distance |
| TITAN [22] | 2014 |  | ✓ |  |  | Two-chain factorial HMM/EM |
| SubcloneSeeker [23] | 2014 | ✓ | ✓ | ✓ |  | Clustering, enumeration and co-localization prediction |
| BTP [24] | 2014 | ✓ |  |  |  | Binary tree partition |
| BreakDown [25] | 2014 |  |  | ✓ |  | Maximum likelihood |
| PhyloSub [26] | 2014 | ✓ |  |  | ✓ | Tree-structured stick-breaking process prior, MCMC |
| BayClone [27] | 2015 | ✓ |  |  | ✓ | Categorical Indian Buffet Process |
| PhyloWGS [5] | 2015 | ✓ | ✓ |  | ✓ | Tree-structured stick-breaking process prior, MCMC |
| CITUP [28] | 2015 | ✓ |  |  | ✓ | Quadratic integer programming |
| LICHeE [29] | 2015 | ✓ |  |  | ✓ | Clustering and evolutionary constraint network |
| AncesTree [30] | 2015 | ✓ |  |  | ✓ | Integer linear programming |
| SPRUCE [6] | 2016 | ✓ | ✓ |  | ✓ | Combinatorial enumeration |
| Canopy [7] | 2016 | ✓ | ✓ |  | ✓ | MCMC |
| THEMIS (our work) | 2017 | ✓ | ✓ |  | ✓ | Dynamic graphical models |

Most existing tools are designed to analyze a single tumor biopsy and are not suitable for jointly analyzing multiple biopsies. As DNA sequencing becomes more affordable, we can more easily collect multiple biopsies from a single patient during treatment. If we only perform heterogeneity analysis on the individual biopsies, then we are unable to detect clones that are shared across different biopsies from the same patient, and we fail to address important questions about how the tumor cells evolve, metastasize and escape treatment.

Finally, although most of models are free and publicly available, it is difficult to extend them by adding new assumptions and new types of biological data. Even under the best of circumstances, significant effort is required for users to fully understand the source code. In many situations, data structures and computational algorithms prohibit other investigators from modifying the model to accommodate their special needs.

To address these challenges, we propose THEMIS (Tumor Heterogeneity Extensible Modeling via an Integrative System), which allows us to jointly characterize different types of genomic aberrations from multiple biopsies using a dynamic graphical model. We implement our model via an extensible modeling platform, the Graphical Models Toolkit (GMTK) [31], which makes our approach open, reproducible and easy for others to extend. To extend the model, users only have to modify the model specification files; GMTK then automatically handles the required computation. Simulation experiments demonstrate that THEMIS significantly increases the accuracy of recovering tumor subclones and their genotypes, compared with its ancestor, TITAN [22]. Single cell DNA sequencing confirms that individual nuclei can be segregated into one of the two tumor subclones identified by THEMIS. We applied THEMIS to three tumor biopsies from one cancer patient, thereby revealing the mutation accumulation history of the patient, tracking cancer progression, and identifying mutations related to developing resistance following various treatments.

## Results

### The Model

From bulk next generation sequencing data, we define two primary observations at each genomic position: the allelic ratio, defined as the proportion of the reads containing a specified allele among all reads aligned to the site, and the log ratio between tumor read depth and normal read depth (**Fig. 1a**). From these inputs, we aim to infer the number of distinct clones, the full genotype of each clone, and the prevalence of each clone within each biopsy (**Fig. 1b**). To carry out this inference, TITAN [22] uses a dynamic graphical model, in which each frame represents one genomic position, and the allelic ratio and tumor/normal log ratio are observed at each frame. The backbone of the TITAN model consists of two hidden Markov chains, one representing the genotype of the CNA event at the current position, and the other representing the clone in which the CNA event occurs. Our model, THEMIS, is similar to TITAN in the sense that both models are dynamic graphical models with each frame representing a single genomic position, with CNA events captured by hidden Markov chains (**Supplementary Fig. 1**). Therefore, THEMIS inherits key assumptions from TITAN (**Supplementary Note 1**). However, THEMIS extends TITAN by (1) jointly accounting for SNVs and CNAs, (2) jointly analyzing multiple biopsies, (3) estimating transition probabilities between hidden states of the model from observed data rather than fixing them at specific values, and (4) using an open and extensible modeling language (GMTK [31]).

**Figure 1:**
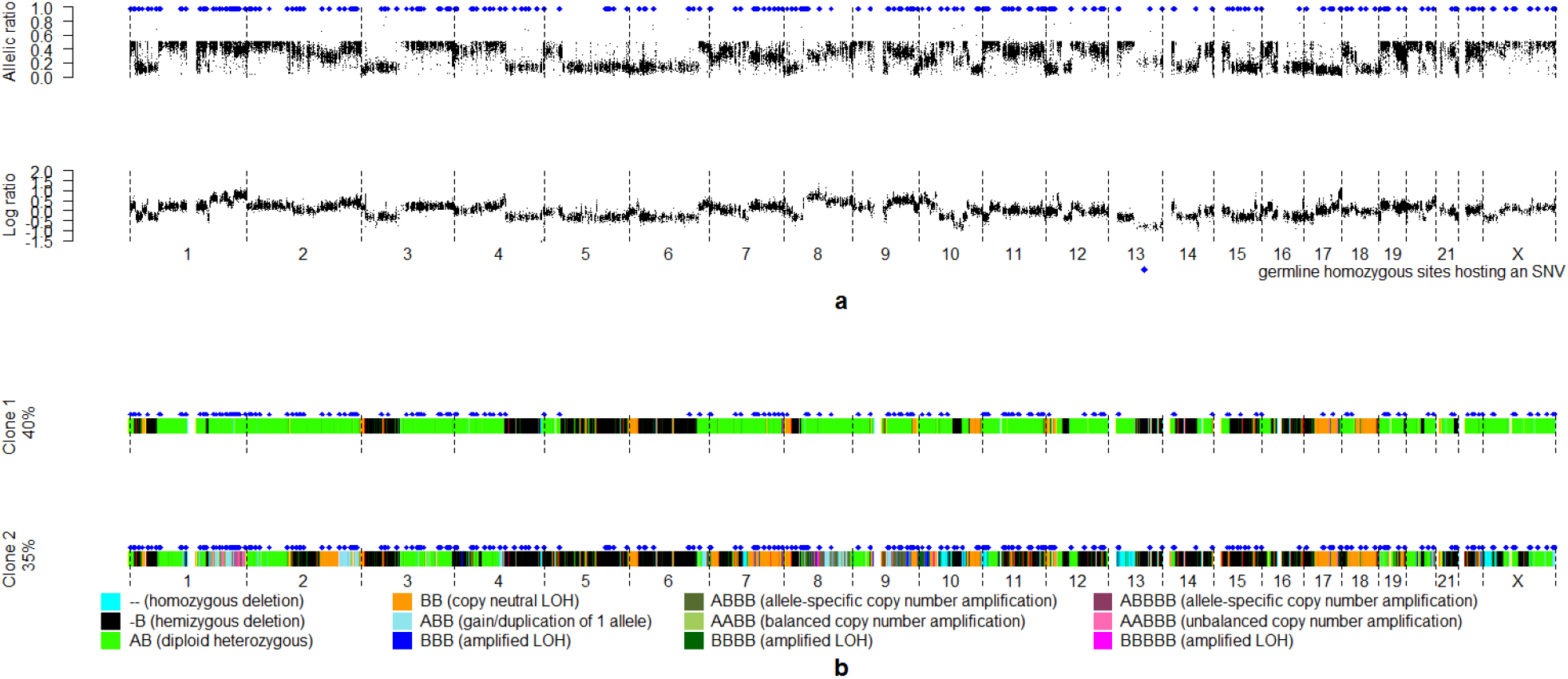
Example THEMIS input observations and the corresponding inferred outputs. (a) Inputs to THEMIS, including allelic ratio, log ratio and genomic position information. Germline homozygous sites are indicated by blue diamonds. (b) Outputs of THEMIS show that there are two tumor clones in the tumor biopsy, one parent tumor clone with 40% cell prevalence and one child tumor clone with 35% cell prevalence. The CNAs of the two tumor clones are visualized with color bars across the genome, and the genomic positions hosting a SNV are indicated by blue diamonds.

### Simulation results

We first used simulated data to compare the performance of THEMIS and TITAN [22]. As a starting point for the simulation, we used the genomic positions measured in three tumor biopsies from three patients with triple negative breast cancer (**Supplementary Table 1**). For each set of genomic positions, we also specified three different sets of tumor subclone compositions. More details about the simulation experiments are provided in **Supplementary Note 2**. We evaluated (1) the percentage of sites at which the hidden genotype was incorrectly inferred, (2) the percentage of sites at which the clonal/subclonal status was incorrectly inferred, and (3) the percentage of sites at which either the genotype or clonal/subclonal status were incorrectly inferred. THEMIS outperformed TITAN in recovering the clonal/subclonal status and genotypes of the genomic positions in all experiments (two-sided paired t-test, *p*=0.00137 for genotype recovery, *p*=1.152 × 10^−7^) for clonal/subclonal status recovery, and *p*=0.00198 for both genotype and clonal/subclonal status recovery), and reduced the recovery error by 13.3% on average (**Supplementary Table 2**). Not surprisingly, both THEMIS and TITAN performed better when the prevalence of the somatic events was higher.

### Validation via single-cell DNA sequencing

The two tumor clones in **Fig. 1b** were identified from the bulk DNA sequencing data from an involved axillary node in a patient with metastatic triple negative breast cancer. We single-cell sequenced a second sample taken from the same axillary node at the same time, aiming to validate the subclones previously identified from bulk DNA sequencing. From a total number of 143,108 nuclei, 96 nuclei were fluorescence activated cell sorted (FACS) using gating to select for tumor cell nuclei, placed on a 96-well plate, and whole genome amplified (GenomiPhi). Indexed Nextera libraries were sequenced on the NextSeq using PE 150 bp mid-output flow cell. Six cells were removed due to extremely low numbers of reads after adapter removal (**Supplementary Fig. 2**). The sequencing data from another eleven nuclei was of low quality, as evidenced by a much larger fraction of short reads (**Supplementary Fig. 3**) and were excluded from our analysis. The remaining 79 cells were used for validation.

**Figure 2:**
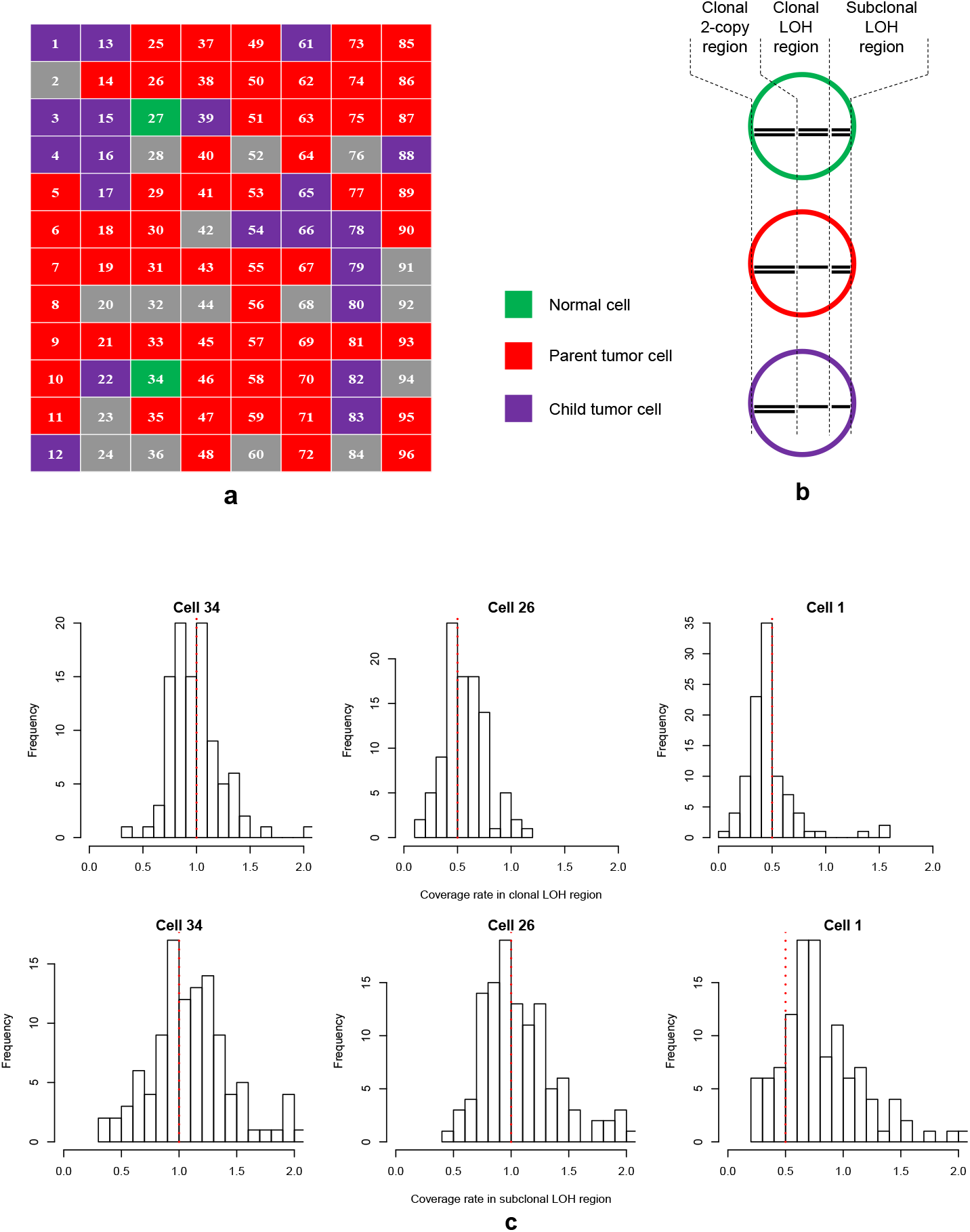
Results of single cell validation experiments. (a) Predicted cell category from the Bayesian classifier: normal cell (green), parent tumor cell (red), child tumor cells (purple) and unknown due to low sequencing quality (grey). (b) Three different types of regions that we use to distinguish the three types of cells, namely clonal 2-copy regions, clonal LOH regions and subclonal LOH regions. (c) Histograms of the relative coverage rate in clonal LOH segments and subclonal LOH segments demonstrate cell 34 is normal, cell 26 is from the parent tumor clone and cell 1 is from the child tumor clone. Red dotted lines in the histograms indicate the expected coverage rates in the cells.

We use a Bayesian classifier (**Supplementary Note 3**) to assign each of the 79 cells into one of the three clones identified from the bulk sequencing data by THEMIS — one normal clone, one parent tumor clone and one child tumor clone. As input to the classifier, we use sequencing coverage on three types of genomic regions which are derived from the inferred genome-wide genotype for the two tumor clones (**Fig. 1b**), namely, clonal 2-copy (i.e. normal) regions, clonal 1-copy (i.e. loss of heterozygosity [LOH]) regions and subclonal 1-copy regions (**Fig. 2b** and **Supplementary Table 3**). Clonal 2-copy regions provide the baseline measurement of sequencing coverage in normal regions. Clonal LOH regions distinguish tumor from normal. Subclonal LOH regions distinguish the child tumor clone from its parent. The Bayesian classifier identifies 2 normal cells (indicating that FACS gating worked), 57 parent tumor cells and 20 child tumor cells (**Fig. 2a** and **Supplementary Table 4**). The observed ratio of parent to child tumor cells (2.85) does not agree with the ratio inferred by THEMIS (1.14), which is likely attributable to the very small number of examined events and that the single cell analysis was performed in a separate sample taken at the same time. However, the histograms of the normalized coverage rate in the 99 clonal LOH segments and 119 subclonal LOH segments in the three types of cells validate that the three categories of nuclei agree with our model (**Fig. 2c**). For example, cell 34 has no LOH events in the clonal LOH region nor in the subclonal LOH region, and is therefore a normal cell. Cell 26 displays LOH events in the clonal LOH region, but not in the subclonal LOH region, and is therefore identified as belonging to the parent tumor clone. Cell 1 has LOH events both in clonal and subclonal LOH regions, and is therefore identified as belonging to the child tumor clone. The histograms of the coverage rates from the aggregated 57 parent tumor cells show similar patterns to that of cell 26, whereas the histograms from the aggregated 20 child tumor cells show similar patterns as cell 1 (**Supplementary Fig. 4**). Therefore, our single cell experiment successfully validates the subclones identified by THEMIS.

### Joint analysis over multiple biopsies from the ITOMIC study

The Intensive Trial of OMics In Cancer (ITOMIC-001) enrolls patients with metastatic triple negative breast cancer in whom biopsies of multiple metastatic sites are performed repeatedly over time [32]. For each patient, multiple biopsies are evaluated using next generation sequencing. We performed joint heterogeneity analysis on three biopsies from Patient 1 (**Fig. 3a**). Patient 1 was originally diagnosed with triple-negative breast cancer in February 2011, and enrolled in ITOMIC-001 in October 2013. The patient then received three different treatments, including cisplatin (between study days 12 and 125), an investigational PARP inhibitor veliparib (between study days 126 and 194), and the kinase inhibitor ponatinib (from study day 195 until the time of her death on study day 250). The first biopsy B1 was sampled from an involved right axillary lymph node, collected on study Day 7 (before cisplatin). The second biopsy B2 was sampled from the same right axillary lymph node on study Day 125 (after cisplatin). The third sample B3 was from a left peribronchial lymph node, collected at autopsy following the patients death on study day 250 (post ponatinib).

**Figure 3:**
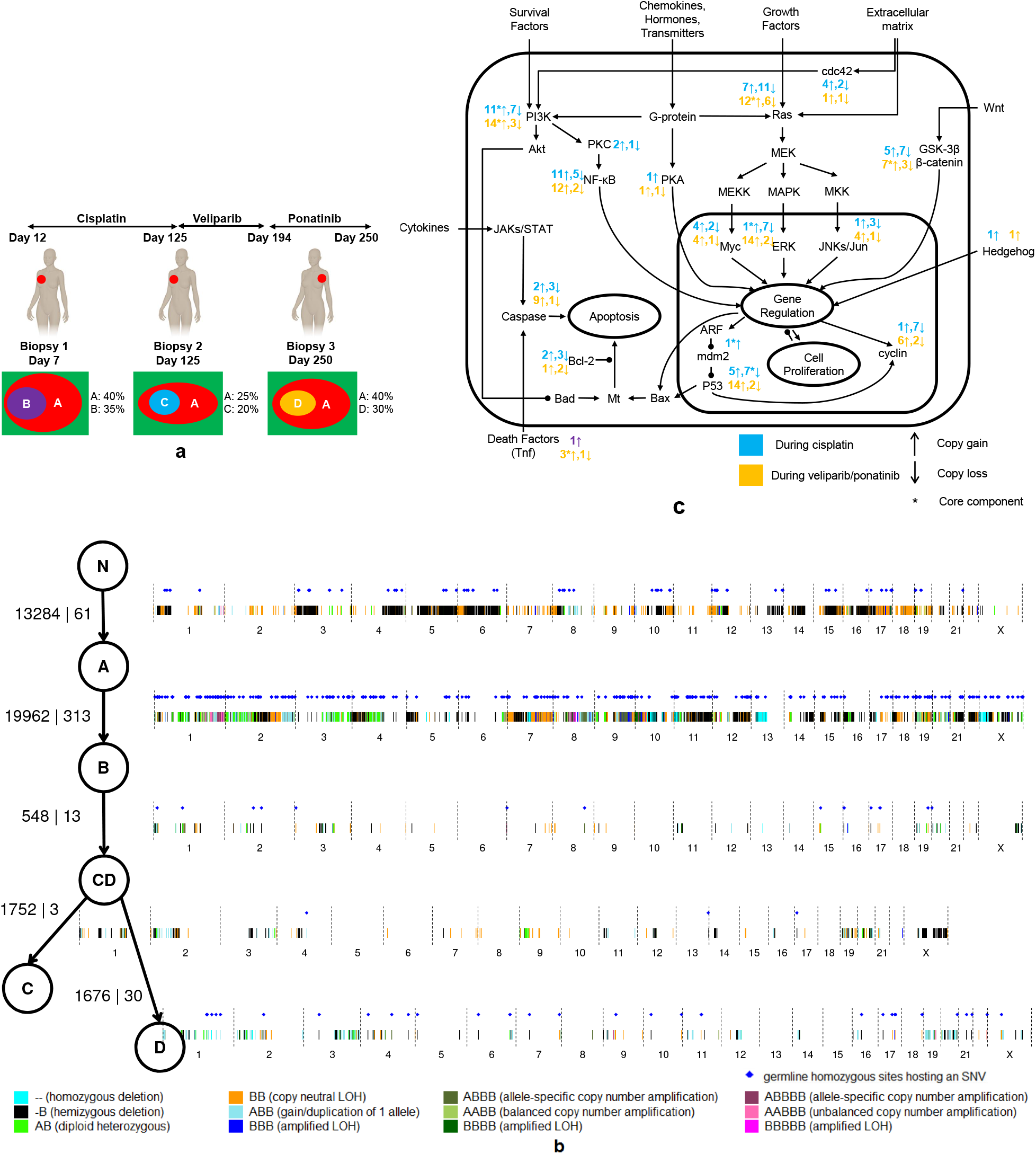
Experiments and results from joint analysis of three biopsies from the same patient. (a) The collection of the three biopsies from the same patient during three stages of treatment and the inferred tumor clones in the biopsies. (b) The recovered phylogenetic tree and the mutations accumulated on each edge. The mutations are shown on a genome-wide plot, and the two numbers on each edge are the number of germline heterozygous sites affected by these CNAs and the number of SNVs. (c) Signal pathways (a simplified version of signal transduction pathways from Wikipedia), and the number of genes with copy number changes (copy gain and copy loss) in different stages of cancer progression. * denotes at least one of the mutated genes is a core component of the signaling pathway.

We followed a five-step procedure to recover the phylogenetic tree from the three tumor biopsies (see Methods for details), which involves enumerating all possible candidate phylogenies from individual biopsy analysis and then selecting the best phylogeny by likelihood (**Supplementary Note 4**). The recovered phylogenetic tree (**Fig. 3b**) reveals the relationships among the recovered clones and the mutations accumulated at each stage of cancer progression. Parent clone A is shared by all three biopsies, and child clones (B, C, and D) inherit the mutations present in parent clone A and acquire new mutations of their own. Although clones B, C and D occur in separate biopsies, clones C and D appear to be descendants of clone B, meaning that the new mutations acquired in later biopsies occurred within the child tumor clones. In addition to clones A, B, C and D, the phylogenetic tree includes an internal node for an inferred intermediate clone (CD) hosting the mutations shared between clone C and clone D, and corresponding to a splitting point between clones C and D in cancer progression. On the phylogenetic tree, we also label the new CNAs and SNVs acquired on the edges. We visualize these mutations on a genome-wide plot, and provide the number of germline heterozygous sites affected by these CNAs and the number of the SNVs (**Fig. 3b**). The mutations on the edges of the phylogenetic tree reveal the mutation accumulation history of this patient and can help in tracking mutations related to treatment resistance. By the time the patient joined the study, there had been two major phases of mutation accumulation, one corresponding to the mutations accumulated in clone A and the other corresponding to mutations accumulated in clone B. Comparing the two phases, more SNVs were acquired in the second phase. After joining the study and receiving further treatments, the patient acquired additional mutations. The mutations on the edges between clone B and clone C were acquired during treatment with cisplatin, whereas those on the edge between clone CD and clone D were acquired during treatment with the PARP inhibitor veliparib (without response) followed by treatment with the kinase inhibitor ponatinib (which did yield a partial response). Because veliparib failed to affect tumor growth, we attribute the changes acquired at the CD → D transition to ponatinib, which was given based on the presence of two activating mutations affecting FGFR2 (S252W;Y375C) (manuscript submitted).

A number of intriguing patterns were revealed when we looked at the mutations acquired during different phases of treatment. First, many of the genes in CNA regions are known to be related to cancer. On the CNAs acquired during the three treatments, including 147 CNAs during the treatment with cisplatin and 98 CNAs during the treatment with veliparib and ponatinib, we identify 848 genes and 519 genes, respectively. We queried these genes on the NCBI gene website (www.ncbi.nlm.nih.gov/gene) and retrieved 186 genes and 175 genes, respectively, that are known to be related to cancer in the literature. The retrieved genes are related to many important cancer signaling pathways (**Fig. 3c**). On some of the cancer signaling pathways (i.e., MEK/MAPK/Erk, PI3K/Akt/mTor, NF-κB, and p53), core genes (i.e., MAP2K7, PIK3C2A, PIK3CD, PIK3CB, TNFSF9, TNFSF14, NRAS, GSK3B, Notch3, TP53TG5 and SNAI1) experienced copy-number changes during different phases of treatment. In addition, the genes mutated during different phases of treatment show patterns that are potentially illustrative of different therapeutic responses. The proportion of genes experiencing copy number gains in later stages (i.e., on the edge CD → D, during the treatments with veliparib and ponatinib) is much higher than that in earlier stage (on the edges B → CD and CD → C, during the treatment with cisplatin), and the proportion difference is more dramatic in important cancer signaling pathways, including MEK/MAPK/Erk, PI3K/Akt/mTor and NF-κB. These genes mutated in the different phases of treatment also showed different functional focuses. According to DAVID [33, 34], the top function clusters among the genes mutated in the earlier stage are rho GTPase activation, growth factor, DNA damage and ErbB signaling. The top function clusters from the genes mutated in the later stage are DNA damage, ras signaling, nucleotide-binding, zinc-finger, neurotrophin signaling, and endocytosis. Third, a number of SNVs occurred on or near the genes known to be related to cancer signaling pathways, which allows us to investigate them together with the genes mutated due to copy number changes. During treatment with cisplatin, one SNV occurred near an intron/extron boundary within BIN1, which is known to interact with the myc oncoprotein as a putative tumor suppressor. During the treatments with veliparib and ponatinib, SNVs occurred on or near LATS1 (a core component of Hippo-YAP pathway), MGMT (related to DNA damage), IL17RB (related to NF-*κ*B signaling) and APCDD1L (related to wnt signaling), all of which are known to be related to breast cancer. Although additional experiments are needed for further validation, THEMIS provides a powerful computational tool to generate hypotheses from multiple biopsy DNA sequencing data.

## Discussion

THEMIS offers a powerful and extensible modeling framework to jointly capture different types of genomic aberrations in the analysis of multiple biopsies. The integration of CNAs and SNVs in the heterogeneity analysis increases the accuracy of clonal inference relative to previous methods that consider only single types of mutations. For example, if we observe an allelic ratio 0.3 at one genomic position, then the cell prevalence of the SNV should be 60% if the corresponding genotype is AB, but the prevalence should be 85.7% if the genotype is AAB. In such cases, methods that fail to jointly consider copy number information and SNVs can be misled. In addition, the integration of multiple types of mutations allows us to understand cancer comprehensively, and to address important questions such as how the different types of mutations cooperate with each other and what roles they play at different stages of cancer progression.

The joint analysis over multiple biopsies from the same patient provides a complete picture of mutation progression in the patient, which may shed light on how tumor cells escape treatment and metastasize. The ability to analyze multiple biopsies jointly will be increasingly important as DNA sequencing costs continue to decrease. The current turnaround time of analyzing the three biopsies with our model, including both data preprocessing and model running, is just a few hours.

Because THEMIS is built using a general purpose graphical models toolkit, the approach is easy to extend to alternative model architectures. In addition, GMTK provides flexible calculation in both estimation and inference, including exact and approximate inference engines. Based on the available computing resources, the user can easily trade memory with running time. Using this modeling and algorithmic flexibility, we plan in future work to extend THEMIS to account for more complex types of mutations, such as chromothripsis and chromoplexy. We also plan to incorporate a more principled phylogeny reconstruction method into THEMIS. Ultimately, THEMIS will provide a testbed for model development by us and others interested in modeling the full complexity of tumor evolution.

**Data availability** The bulk DNA sequencing data and the single cell DNA sequencing data used in our analysis can be downloaded from Sequence Read Archive with accession SRP102304.

**Software availability** THEMIS is available at https://github.com/jieliu6/THEMIS.

## Methods

### Data preprocessing flow

We assume that next generation sequencing data was mapped to the reference genome, and the mapped BAM files are ready for analysis. There are three major data preprocessing components, including (1) identifying the genomic sites that will be included in the model, (2) screening unreliable sites and reads with MuTect [35], and (3) adjusting GC content and mappability with HMMcopy [36] (**Supplementary Fig. 5**). Our model captures both CNA events and SNV events; therefore, two types of genomic sites are included. For CNAs, we consider *germline heterozygous sites* since we can monitor not only the absolute copy number changes (by looking into tumor-normal read depth difference), but also what happens to the two individual copies (by looking into allelic imbalance). For SNVs, we consider the *germline homozygous sites* which host an SNV event in any of the biopsies. From the germline (normal) BAM file, we use Samtools to identify germline heterozygous sites. From the tumor and normal BAM files, we use MuTect [35] to identify germline homozygous sites which host an SNV. There are still many artifacts in BAM files which could potentially introduce errors, and we remove sites and reads that are not reliable by MuTect [35]. Short reads from next generation sequencers are not uniformly distributed across the genome — more reads are expected to be obtained from regions with higher GC content and mappability. The bias cannot fully be adjusted by normalizing with another next generation sequencing library (e.g. from a normal biopsy) from the same patient [37]. We therefore use HMMcopy [36] to adjust GC content and mappability in the read counts.

### Structure of the THEMIS model

In the graphical model representation in **Supplementary Fig. 1**, each vertex represents one random variable at a particular genomic position. If a vertex is shaded, then the corresponding random variable is *observed*. Otherwise, the random variable is *hidden* and hence unshaded. The variables and parameters used in the model are listed in **Supplementary Table 5**. We use capital letters to denote random variables and the corresponding lower-case letters to denote the particular values of the random variables. If a lower case letter has a bar on top, then it is observed. Otherwise, it is inferred. There are two hidden chains, i.e. two hidden variables at each site *t*, corresponding to the unknown genotype of the mutation (*G_t_*) and in which clone the mutation occur (*Z_t_*), respectively. At site *t* and in biopsy *m*, a set of observed variables reflect the allelic ratio (*A_m,t_*), log ratio between tumor read depth and normal read depth (*L_m,t_*) and the type of site (*D_t_*). In biopsy *m*, there are another set of |*Z*| hidden variables, 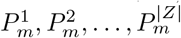, denoting the prevalence levels of the clones. Between any two adjacent sites, we specify the *transition probability* between genotypes and the transition probability between clones. At each site, we also specify the *emission probability* of the observed variables given the hidden variables. At a germline heterozygous site (where *D_t_*=0), the allelic ratio reflects how the allelic ratio is different from 0.5, which is expected in a normal cell. At a germline homozygous site (where *D_t_*=1), the allelic ratio reflects how the allelic ratio is different from 0, which is expected in a normal cell. Therefore, the parents of the allelic ratio (*A_m,t_*) include the genotype (*G_t_*), the clone index (*Z_t_*), the prevalence levels 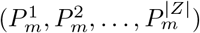 and the type of site (*D_t_*). Since the log ratio of tumor-normal read depth difference (*L_m,t_*) does not depend on the type of site, its parents include the genotype (*G_t_*), the clone index (*Z_t_*) and the prevalence levels 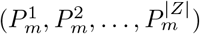. The binary variable *S_t_* denotes the start of a chromosome. When the current site is the start of a chromosome (*S_t_*=1), *G_t_* and *Z_t_* do not depend on *G_t_*_−1_ and *Z_t_*_−1_, but follow uniform prior distributions. The variable *H_t_* is the distance (in base pairs) between site *t* and its previous site *t* − 1. We set transition parameters (for both *G_t_* and *Z_t_*) as functions of 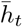 to capture the phenomenon that the chance *G_t_* and *G_t_*_−1_ agree decreases as 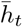 increases (same for *Z_t_* and *Z_t_*_−1_). Therefore, *H_t_* is also a parent of *G_t_* and *Z_t_* at a non-start-of-a-chromosome site *t*.

### Variables in the THEMIS model

The genotype *G_t_* at site *t* is a discrete variable, which corresponds to all possible genotypes up to a certain number of copies. We consider all possible genotypes up to 5 copies (**Supplementary Table 6**). The clone index variable *Z_t_* at site *t* is a discrete variable of |*Z*| possible values (i.e., *Z_t_* ∈ {1, …, |*Z*|}), where |*Z*| is pre-specified by the user. For clone *z* in biopsy *m*, the prevalence level variable 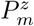 is a discrete variable of |*P*| possible values, where |*P*| is pre-specified by the user. The default |*P*| is 20, corresponding to the 20 equally spaced prevalence levels between 0 and 1 (i.e., 0.05, 0.10, …, and 1.00). The site type *D_t_* at site *t* is a discrete variable with *D_t_*=0 denoting a germline heterozygous site and *D_t_*=1 denoting a germline homozygous site which hosts a SNV event. The allelic ratio *A_m,t_* at site *t* in biopsy *m* is modeled as a Gaussian variable, i.e.,

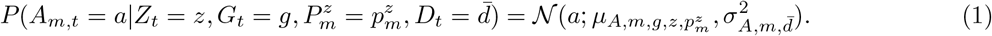

The mean parameter 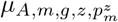 is

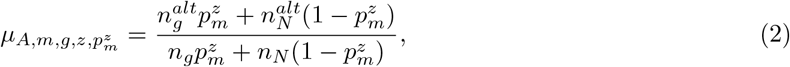

where *n*_*g*_ is the DNA copy number in tumor cells with genotype *g, n*_*N*_ is the DNA copy number in normal cells, 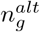 is the copy number of the alternative allele in tumor cells with genotype *g*, and 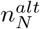 is the copy number of the alternative allele in normal cells (**Supplementary Table 6**). The mean parameter 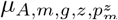 is not estimated from data, but determined by the states of the hidden variables *G_t_, Z_t_*, and 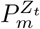 and the observed 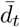. The variance parameter 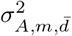 is estimated from data. The log ratio of tumor-normal read depth at site *t* in biopsy *m*, denoted by *L_m,t_*, is modeled as a Gaussian variable, i.e.,

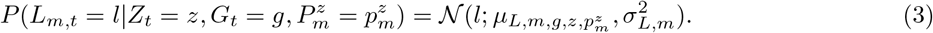

The mean parameter 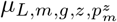 is 
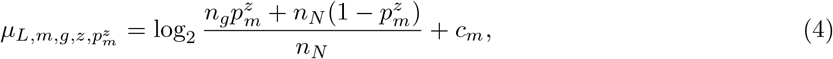
where *n_N_* is copy number in normal cells (set to be 2 by default), and *n_g_* is the DNA copy number in tumor cells with genotype *g*. The parameter *c_m_* captures the sequencing depth difference in the tumor biopsy and the normal biopsy and the read number discrepancy due to ploidy change in the tumor biopsy. Therefore, the mean parameter 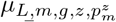 is not estimated from data, but determined by the states of the hidden variables *G_t_, Z_t_*, and 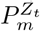. The variance parameter 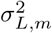 is estimated from data.

### Parameters in the THEMIS model

There are three sets of parameters that are estimated from data. The first set of parameters include the variance of allelic ratio in biopsy *m* on germline heterozygous sites, denoted by 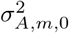, the variance of allelic ratio in biopsy *m* on germline homozygous sites, denoted by 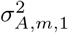, and the variance of log ratio in biopsy *m* on each site, denoted by 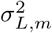. The second set of parameters include the transition probability from genotype *j* at site *t* − 1 to genotype *i* at site *t* (*i*; *j* ∈ {1; …, |*G*|), denoted by 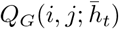 and transition probability from clone *j* at site *t* − 1 to clone *i* at site *t* (*I, j* ∈ {1; …, |*Z*|), denoted by 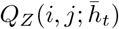. We model 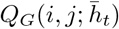 and 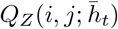 as parametric functions of 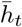, the distance (in base pairs) between site *t* and site *t* − 1. We first define the probability of staying at the same genotype *j*,denoted by 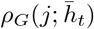 as

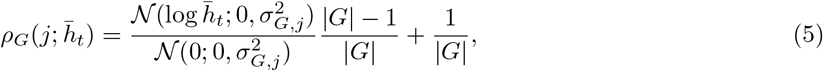

where 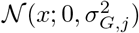 is the probability density of a Gaussian distribution with mean 0 and variance 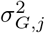 at the point *x*. Then

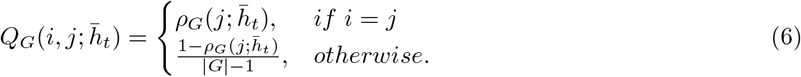

Similarly, we define

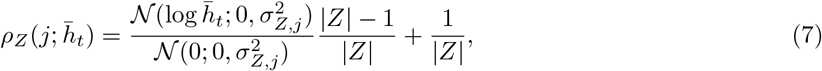

and

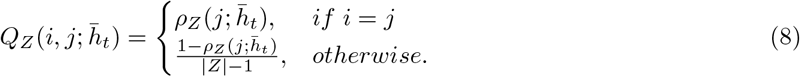

Note that we estimate 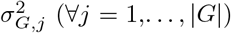 and 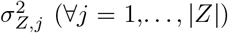 from data in a maximum likelihood fashion. The way we parameterize the transition probabilities captures the phenomenon that the probability of staying at the same genotype decreases as a monotone function of the distance from the previous site, and the rate it decreases is parametrized by 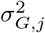. Therefore, we estimate the decreasing speed (i.e. 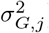) adaptively from the data (unlike user-prespecified transition probabilities in TITAN), and the speed is different for different genotypes since different mutation events may occur with different lengths on the genome (unlike the tied transition probabilities in TITAN). The third set of parameters include the prior distributions of genotypes, clones and the cell prevalence levels of the clones (denoted by π_*G*_, π_*Z*_ and π_*P*_). These prior distributions are responsible for the frames that correspond to the starts of the chromosomes. These prior distributions are initialized as uniform distributions, and trained from the data along with other parameters. Therefore in Formula (9), the parameter set 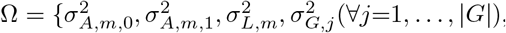, 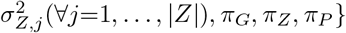

The user has to decide and specify the following parameters, the number of biopsies used in the analysis (denoted by *M*), the number of subclones (denoted by |*Z*|), the maximum copy number in the mutations (denoted by 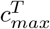), the number of prevalence levels (denoted by |*P*|), and log-ratio offset in biopsy *m* due to ploidy and sequencing depth change (denoted by *c_m_*).

We use Ω to denote the set of parameters in the joint distribution specified by the model. In the estimation step, we find a (local) maximum, denoted by 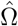, for

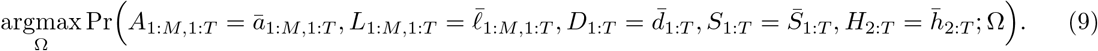

In the inference step, we infer the most probable states of the hidden variables given the estimated parameters 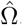, i.e.,

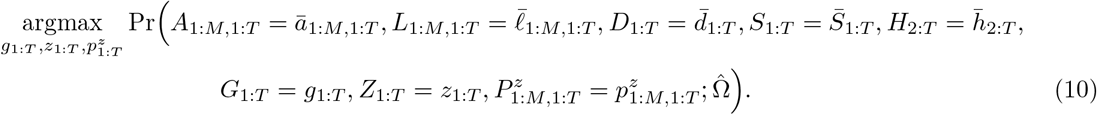

The inferred most probable sequence of hidden variables, as the output of our algorithm, provide us the heterogeneity analysis results, i.e., a number of subclones and their genotypes and cell prevalences (**Fig. 1b**).

### Selecting the number of subclones

THEMIS requires the user to specify the number of subclones in the biopsies before running the model. There are three ways of identifying the number of subclones from the data. The first method is a naive eyeball method. If the biopsy is well-sequenced, the number of subclones can be directly identified from the bivariate plot (allelic ratio against log ratio at germline heterozygous sites) of the biopsy by observing different prevalence levels of the LOH events. We take one tumor biopsy (biopsy B1 in ITOMIC study) as an example, whose bivariate plot is provided in **Supplementary Fig. 6**. It can be observed that there are two major LOH prevalence levels in the plot. Therefore, we can assume that there are two tumor subclones in the biopsy. Another way of choosing the number of subclones is to use the Bayesian information criterion (BIC) [38]. BIC is defined as 
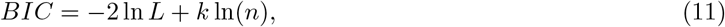

where ln*L* is the log likelihood of the data, *k* is the degree of freedom, and *n* is the number of data points. We choose the subclone number which produces the smallest BIC score. When we run the 2-subclone model and the 3-subclone model on the tumor biopsy B1, BIC scores are −476,725 and −464,710, respectively. Therefore, we can assume there are two tumor subclones in the biopsy based on the BIC scores. Another way of choosing the number of subclones is to use cross-validation (CV). Suppose that we use three-fold cross-validation. We randomly partition the chromosomes into three sets. In each training-testing split, we use the data from two sets as the training data and the remaining set as the testing data. With the estimated parameters from training data, we run Viterbi algorithm on the testing data, and choose the subclone number which produces the largest averaged log-likelihood (a.k.a., the Viterbi score in GMTK) on testing data. In both simulated data (in Simulations 1 and 2, we simulated two tumor subclones and three tumor subclones, respectively) and real data (biopsy B1 in ITOMIC study), we observed that the three methods provide the correct results (**Supplementary Table 7**). In practice, users may use any of the three methods or a combination of the three methods to set the number of subclones. Note that although the naive eyeball method is straightforward, it may produce inaccurate estimate if the sequencing depth of the biopsy is low or when the prevalence levels of two subclones are close to each other. CV is more robust compared with BIC, but requires additional computation cost. When BIC and CV are being used, we recommend the users to start with a small number (e.g. 2 subclones) and increase the number until the evaluation criteria deteriorate. For example, if 2-subclone model produces lower BIC (or higher averaged ln*L* in CV) than 3-subclone model, they no longer have to run 4-subclone model.

### Joint analysis multiple biopsies from the same patient

In addition to the assumptions we make for individual biopsy analysis (**Supplementary Note 1**), we make three additional assumptions for joint analysis over multiple biopsies. First, we assume that we have the statistical power to discern all the clones from individual biopsies and estimate their prevalences. Second, we assume that we can identify shared clones between biopsies by computing and thresholding similarities between the clones. Third, if the sum of the prevalences (*p_a_* and *p_b_*) of clones a and b is greater than 1.0 in at least one biopsy, and *p_a_* > *p_b_* in all biopsies where clones *a* and *b* are present, then clone *a* is an ancestor of clone *b*.

The joint analysis over multiple biopsies can be carried out in the following five steps. Step 1. Analyze biopsies separately with THEMIS and identify the genotype and prevalence of each clone within each biopsy. Step 2. Compute similarity between all pairs of clones from different biopsies and merge similar clones. Step 3. Identify consistent parent-child relationships based on the individually estimated prevalences using the third assumption above. Step 4. Enumerate all phylogenies consistent with those relationships. Step 5. Select the phylogeny with maximum likelihood.

## Acknowledgments

The authors would like to thank the patient and her family. We also gratefully acknowledge the support from the Washington Research Foundation Fund for Innovation in Data-Intensive Discovery, the Moore/Sloan Data Science Environments Project at the University of Washington, and South Sound CARE.

## Author contributions

J.L., J.A.B. and W.S.N. conceived the method. J.L. and J.T.H. performed the analysis. J.L., W.S.N, and C.A.B. wrote the manuscript. E.M.M., C.S., M.O.D., and C.A.B. contributed experimental results. S.B., V.K.G., and C.A.B. contributed to the clinical trial. R.M.D., C.L., E.M.M., D.P., P.R., and J.S. acquired single cell sequencing data.

